# A reliable and efficient adaptive Bayesian method to assess static lower limb position sense

**DOI:** 10.1101/2023.01.23.525102

**Authors:** Jonathan M Wood, Susanne M Morton, Hyosub E Kim

## Abstract

**Background:** Lower limb proprioception is critical for maintaining stability during gait and may impact how individuals modify their movements in response to changes in the environment and body state, a process termed “sensorimotor adaptation”. However, the connection between lower limb proprioception and sensorimotor adaptation during human gait has not been established. We suspect this gap is due in part to the lack of reliable, efficient methods to assess global lower limb proprioception in an ecologically valid context.

**New Method:** We assessed static lower limb proprioception using an alternative forced choice task, administered twice to determine test-retest reliability. Participants stood on a dual-belt treadmill which passively moved one limb to stimulus locations selected by a Bayesian adaptive algorithm. At the stimulus locations, participants judged relative foot positions and the algorithm estimated the point of subjective equality (PSE) and the uncertainty of lower limb proprioception.

**Results:** Using the Bland-Altman method, combined with Bayesian statistics, we found that both the PSE and uncertainty estimates had good reliability.

**Comparison with Existing Method(s):** Current methods assessing static lower limb proprioception do so within a single joint, in non-weight bearing positions, and rely heavily on memory. One exception assessed static lower limb proprioception in standing but did not measure reliability and contained confounds impacting participants’ judgments, which we experimentally controlled here.

**Conclusions:** This efficient and reliable method assessing lower limb proprioception will aid future mechanistic understanding of locomotor adaptation and serve as a useful tool for basic and clinical researchers studying balance and falls.

## 1. Introduction

Lower limb proprioception is critical for maintaining upright stability and regulating the gait cycle. The proprioceptive sense is formed through a combination of inputs, mostly from muscle spindles and Golgi tendon organs, but also cutaneous and joint capsule receptors (Proske and Gandevia, 2012). Here, we are most interested in proprioception as it relates to static lower limb position sense. Static lower limb position sense contributes to balance and stability as evidenced by studies showing that impaired static lower limb proprioception is associated with higher fall risk (Lord et al., 1991; Lord and Ward, 1994; Ribeiro and Oliveira, 2007). In addition to its contributions to posture and stability, static lower limb proprioception may also impact how individuals implicitly adapt their gait pattern in response to changes in the environment or body state (e.g., fatigue) (Bruijn et al., 2012; Bunday and Bronstein, 2009), a learning process termed “sensorimotor adaptation” (Prokop et al., 1995; Reisman et al., 2005). This link between static proprioception and sensorimotor adaptation is well established in upper-extremity reaching (Cressman and Henriques, 2009; Harris, 1963; Henriques and Cressman, 2012; Mattar et al., 2013; Ostry et al., 2010; Simani et al., 2007; Tsay et al., 2022). However, no causal link between proprioception and locomotor adaptation has been clearly established in humans, which is surprising given how dependent normal walking is on reliable proprioceptive estimates (Dietz, 2002; Hiebert et al., 1996; Kriellaars et al., 1994; Pearson, 2004; Roden-Reynolds et al., 2015; Whelan et al., 1995). We suggest that this discrepancy exists because of the way proprioception is measured in the lower limb. Specifically, most lower limb proprioception assessments are not performed in an ecologically valid context, and those that are lack established reliability and validity (Sombric et al., 2019; Vazquez et al., 2015). In order to address these limitations, we have developed a new psychophysical method for assessing lower limb proprioception.

We first considered the measurement of static lower limb proprioception in a context that most closely approximates gait. A static test allowed us to isolate, as best as possible, the contribution from lower limb proprioceptors to position sense, without help from an efference copy associated with voluntary movement (Wolpert et al., 1995). Furthermore, the importance of static proprioception to gait has been shown in studies assessing falls risk. Current methods assessing static lower limb proprioception often do so within a single joint, in a non-weight bearing position, and rely heavily on remembered positions (Han et al., 2016; Hillier et al., 2015; Horváth et al., 2022). While these methods have proven useful for characterizing specific deficits in joint proprioception after orthopedic injury (e.g., Relph et al., 2014), they cannot be readily translated to functional lower extremity movements. This is because a unified percept of limb location results from the nervous system’s integration of proprioceptive signals across multiple limb joints (Bosco et al., 2000; Fuentes and Bastian, 2010; Gandevia, 1985; Proske and Gandevia, 2012; Soechting, 1982). The body position in which proprioception is measured is also important because differences in proprioceptive estimates emerge in weight bearing vs non-weight bearing positions both in the upper limb (Ansems et al., 2006) and in the knee joint (Bullock-Saxton et al., 2001; Stillman and McMeeken, 2001). For these reasons, we measured whole lower limb proprioception while standing as this provides the closest approximation to gait.

A comprehensive way of characterizing a sensory system is using a psychophysical assessment to estimate two of its distinct physiologic characteristics: the point of subjective equality (PSE) and the uncertainty. The PSE is the stimulus that is perceived as equal to an internal standard. In this case, the internal standard is symmetry of the left and right limbs. Uncertainty refers to the variability in responses surrounding the PSE, reflecting the noise within the sensory system. The PSE and uncertainty seem to play distinct roles in sensorimotor adaptation (Ruttle et al., 2021; Tsay et al., 2021). While the upper extremity PSE shifts as a result of sensorimotor adaptation, proprioceptive uncertainty at baseline predicts the magnitude of sensorimotor adaptation. The proprioceptive shift and uncertainty estimates themselves are uncorrelated across individuals (Tsay et al., 2021). Thus, it is important to reliably estimate both the PSE and uncertainty.

A widely accepted method of estimating both the PSE and uncertainty is to use a two-alternative forced choice (2AFC) task, where, in the context of the current study, participants judge the relative positions of their feet when placed at various locations. The response data are then fit with a psychometric function, where the inflection point (i.e., the probability of a response being 0.5), represents the PSE and the slope of the function is inversely proportional to the uncertainty (Kingdom and Prins, 2016). We know of two studies that used a 2AFC task to estimate both PSE and uncertainty of lower limb proprioception (Vazquez et al., 2015; Waddington and Adams, 1999). However, the Waddington study used active repositioning focused on only ankle inversion and eversion movements, whereas the Vazquez study did not measure or report the reliability of their method. Furthermore, in the latter study participants’ responses were heavily impacted by the direction in which the test limb was moved to the stimulus position, which was either always forward or always backward depending on group assignment. Ideally, in a static assessment of lower limb proprioception, the movement of the limbs to stimulus positions should not exert any influence on participants’ judgments. Therefore, while a method like the one used in the study by Vazquez and colleagues offers promise, adjustments are needed to improve the validity of responses, and the reliability of the estimates must be assessed.

Efficiency is also critical for assessing changes in proprioception in single-session motor learning studies, when considering factors such as clinical feasibility, and to maintain participant safety, as standing still for long periods can lead to syncope (Jardine et al., 2018). In the most established psychophysical method, the method of constant stimuli, participants make the same number of sensory judgements at each pre-determined stimulus location. However, this method is inefficient because some of the stimulus locations provide little information about the PSE or uncertainty estimates (Kingdom and Prins, 2016; Leek, 2001; Watson and Fitzhugh, 1990). Indeed, Kingdom and Prins (Kingdom and Prins, 2016, p. 57) suggested that as many as 400 trials are required to accurately estimate both the PSE and uncertainty using the method of constant stimuli. Adaptive psychophysical methods, where the stimulus locations are selected based on prior responses, were specifically developed to solve this efficiency problem (Leek, 2001). The Psi algorithm is one such method that uses Bayesian estimation to calculate PSE and uncertainty values after each trial (Kontsevich and Tyler, 1999). It then chooses the next stimulus that will maximize the information gained for both estimates on the subsequent trial (Kontsevich and Tyler, 1999; Lesmes et al., 2010, 2006; Prins, 2013). This method results in more efficient and accurate estimates of both the PSE and uncertainty compared to the method of constant stimuli (King-smith and Rose, 1997; Kontsevich and Tyler, 1999; Livesey and Livesey, 2016; Turpin et al., 2010). The Psi algorithm is most frequently used in visual and auditory psychophysics, and we note only one instance where is was used to estimate (wrist) proprioception (Elangovan et al., 2018).

Here, we implemented an adapted version of the Psi algorithm and measured its reliability in assessing lower limb proprioception in an ecologically valid context using a test-retest design. We first wanted to ensure that participant responses were not confounded by the direction of movement to each stimulus location, a concern we had based on a prior study (Vazquez et al., 2015). Then, to assess reliability across test sessions, we calculated agreement, defined as the ability for a test to reproduce the same values when measured at different times, using the Bland-Altman method (Altman and Bland, 1983; Giavarina, 2015). In addition, we used Bayesian statistics to fully quantify the probability that the method has good agreement, defined as low evidence of bias in the PSE and uncertainty estimates across the two tests. Combined, we found that, with the current method, movement direction did not impact participant responses, and good agreement for both the PSE and uncertainty could be achieved after only 50 trials.

## 2. Materials and methods

### 2.1. Participants

Young healthy participants between the ages of 18 and 35 were recruited from the University of Delaware community. Participants were excluded if they had any chronic or recent musculoskeletal or neurologic diagnoses, pain, or impaired sensation. This work was completed in accordance with the Code of Ethics of the World Medical Association. All participants provided written informed consent prior to being enrolled into the study. This study was approved by the University of Delaware Institutional Review Board.

### 2.2. Motion capture

Participants stood on a dual-belt treadmill instrumented with two force plates, one under each belt (Bertec, Columbus OH). We obtained kinematic data, sampled at 100Hz, using an eight-camera Vicon MX40 motion capture system with Nexus software (Vicon Motion Systems Inc., London, UK). Seven retroreflective markers were placed on participants’ shoes and ankles in the following locations: bilateral heels, bilateral lateral malleoli, bilateral 5^th^ metatarsal head, and the right 1^st^ metatarsal head. We used custom written MATLAB scripts (version 2022a, MathWorks, Natick, MA) to control the treadmill belts and obtain live kinematic and kinetic data from Nexus software.

### 2.3. 2AFC task

We used a 2AFC task to measure lower limb proprioception. To measure test-retest reliability, each participant performed the test twice on the same day, with a 20-minute break between tests. For each test, participants stood on a treadmill with vision of their legs occluded with a black drape, and auditory feedback (sounds of the treadmill belts/motors) reduced with noise cancelling headphones (Figure 1A). The primary kinematic variable for the test was foot position difference (in millimeters):

**Figure 1.**
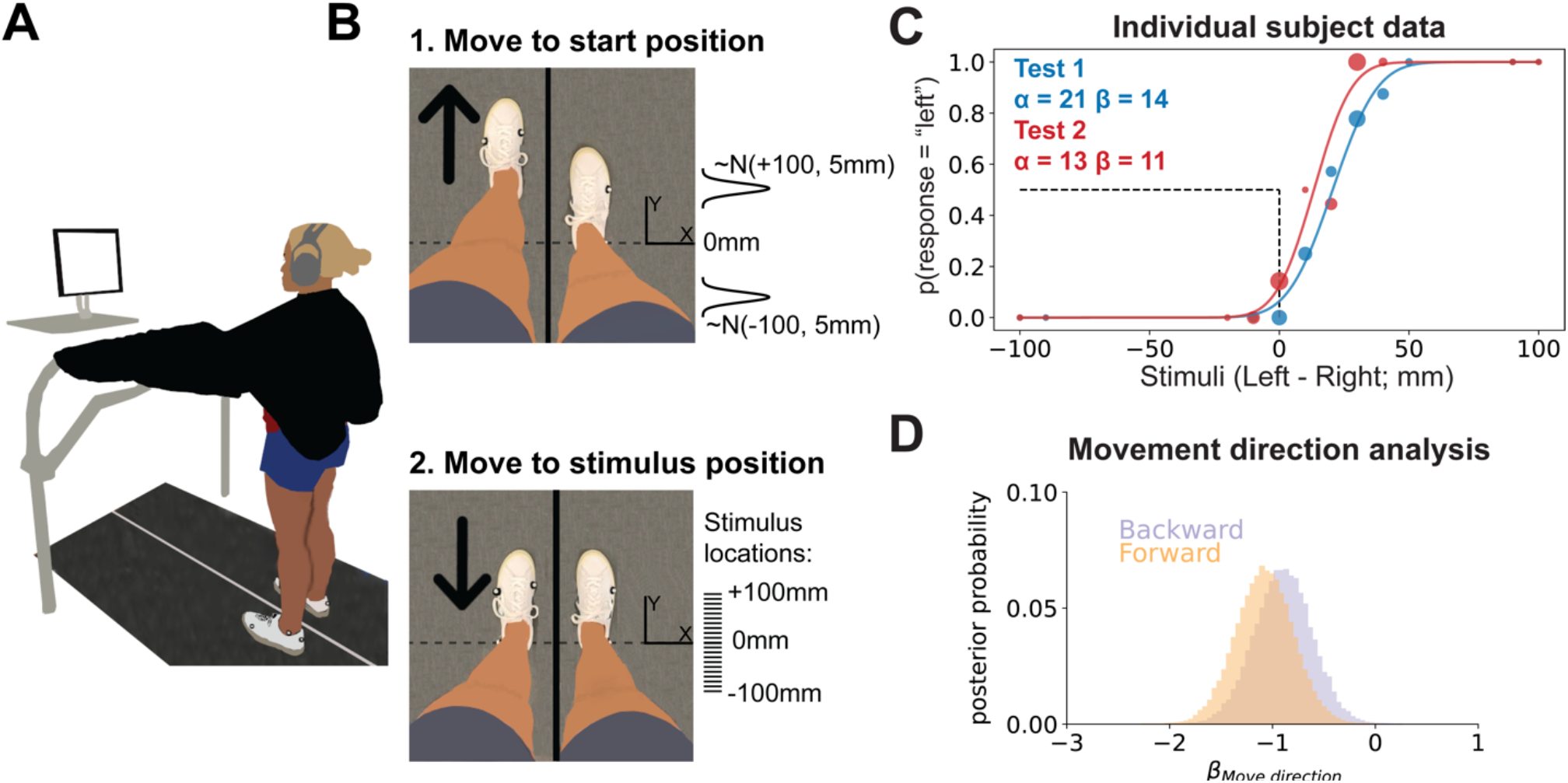
Task setup, representative data, and movement direction analysis. **(A)** 2AFC task setup. Participants stood on the split-belt treadmill with visual feedback of their legs occluded by a black drape and auditory feedback occluded by noise cancelling headphones. The screen prompted participants to verbally respond to the question “Do you feel like your right or left foot is more forward?” when the test foot stopped at the stimulus location. **(B)** Trial sequence. 1) The test foot was moved to a start position either in front of or behind the stimulus position. Start positions were sampled from one of two normal distributions, depending on the movement direction assigned for that specific trial. 2) The test foot was moved to one of 21 possible stimulus positions between -100 and +100 mm. Tick marks (drawn to scale) represent each possible stimulus location. **(C)** Individual participant data. We plotted the psychometric functions that correspond to the estimated PSE (α) and uncertainty (β) values for Test 1 (blue) and Test 2 (red). The dashed black lines, provided as a reference, represent a PSE of 0 mm. The individual dots represent the participant’s response data for each test. Each dot’s size is proportional to the number of trials at that stimulus level. **(D)** Movement direction analysis. To determine if movement direction biased responses, we performed a logistic regression (equations 8a&b) with predictors for subject, stimulus position, and movement direction. Posterior samples of beta coefficients representing contributions of forward vs backward movements to participants’ judgments (β_Move direction_) are plotted as histograms. The large amount of overlap between these distributions indicates that responses were not biased by movement direction (β_Move direction [Backward]_ - β_Move direction [Forward]_ mean [95% HDI] = 0.18 [-0.07 0.43], 99.5% in ROPE).

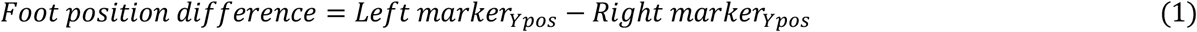

We used the lateral malleoli markers to measure foot position difference. Thus, positive values indicate the left foot was forward of the right foot, while negative values indicate the left foot was behind. Since the foot is the end-effector for the lower limb, we assume judgements were made by combining proprioceptive information across lower limb joints. The right foot served as the reference foot and did not move throughout the test. The left foot served as the test foot and was passively moved by the left treadmill belt to the stimulus positions, measured in terms of foot position difference. After participants performed two practice trials to orient them to the task, their heels were aligned so that everyone started from the same position relative to the laboratory’s y-axis. Since there may still be differences in the ankle markers along the y-axis in this position, we corrected for this baseline difference (rounded to the nearest millimeter) for stimulus positions and PSE estimates. Each test was comprised of 75 trials, and each trial had two parts: 1) movement to a start position, 2) movement to a stimulus position.

To prevent participants from using potential cues other than the stimulus position to determine which foot was more forward, we varied the treadmill speed and distance from the start to the stimulus position on each trial. The treadmill moved the test foot to a start position at a speed selected from a uniform distribution between 40 and 50 mm/s (Figure 1B, top). We controlled for the potential bias in responses caused by movement direction by providing pseudorandomized start positions so that the test foot started in front of the stimulus position 38 times and behind the stimulus 37 times. Therefore, we assume any bias in responses from always moving to the stimulus from the same direction would be washed out, something we confirmed in our formal analysis (see Section 3.1). The specific start position on a given trial, was selected from one of two normal distributions centered around either -100 or +100 mm of foot position difference depending on the movement direction for that trial. Both distributions had a standard deviation of 5 mm:

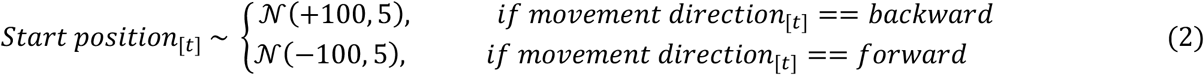

If the sample resulted in the wrong movement direction (e.g., +80 start position for a +90 stimulus on a backwards trial), we resampled from the same distribution until a start position that would result in the correct movement direction was obtained.

After a 0-2 second pause at the start position, the test foot was moved to a stimulus position at a speed selected from a uniform distribution between 10 and 30 mm/s (Figure 1B, bottom). The stimulus positions were located at 21 possible foot difference locations: every 10 mm interval between -100 to +100 mm. Most stimulus positions were selected using the Psi algorithm (see section 2.5). However, we inserted pre-selected stimulus locations to keep participants engaged in the task. There were two types of pre-selected stimuli: 1) far stimuli (±100 or ±90) inserted randomly once every 10 trials, and 2) near stimuli (±10, ±20 or ±30 mm from the current PSE estimate, rounded to the nearest stimulus location) inserted randomly once every 5 trials. No preselected stimuli were inserted within the first 5 trials.

Each time the test foot reached its stimulus position, the treadmill stopped, and a prompt appeared on a monitor in front of the participant: “Do you feel your right or left foot is more forward?” The participant’s response was recorded by the experimenter using a custom graphical user interface (GUI) in MATLAB.

Before the test, participants were instructed to keep their weight equally distributed between both feet, measured with the force plates under each treadmill belt and displayed for the experimenter. If participants consistently kept greater than ∼60% of their weight through one foot, a verbal reminder was provided. We implemented short breaks after 25 and 50 trials to prevent fatigue and blood pooling in the legs. Before the break, current marker positions were recorded from Nexus software, then the participant was asked to walk around the lab for ∼30 seconds. After the participant stepped back onto the treadmill, we positioned their feet so the ankle markers were back to the exact location from which they started and then testing continued.

### 2.4. Psychometric function

Each participant’s probability of responding that the test foot was more forward at each stimulus position (s) was represented as a normal cumulative distribution function (cdf) with two parameters:

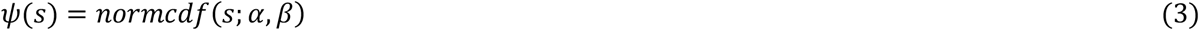

The α parameter, the mean or inflection point of the cdf, represents the PSE, corresponding to the position that the participant perceives the feet were in the same location along the laboratory’s y-axis (i.e., where the probability of judging “left” is exactly 0.5). The β parameter, the standard deviation of the cdf, represents the uncertainty, which reflects the noise within the sensory system itself. Both are measured in terms of foot position difference. The α and β parameters were estimated adaptively on each trial using the Psi algorithm. Of note, in our pilot testing the probability of responding left at -100 and +100 was consistently 0 or 1, respectively. Thus, we did not include the lapse and guess rate parameters that are sometimes estimated in other adaptive algorithms (Prins, 2013).

### 2.5. Psi Algorithm

Here we provide a brief description of the Psi algorithm, which is described in detail by Kontsevich and Tyler (1999). There are two primary components of the Psi algorithm: 1) stimulus selection and 2) parameter estimation.

#### 2.5.1. Stimulus selection

Before each trial (t), the Psi algorithm computes the joint probability of α and β values, given both a “left” and a “right” response (r) at each possible stimulus location (s) for the next trial (*p*_*t*+1_(*α, β*|*s, r*_*t*+1_1). Information entropy (H), a measure of the magnitude of uncertainty in a probability distribution, of the joint posterior distribution is then calculated for both a left and right response:

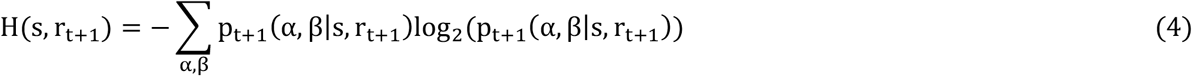

Then, the expected reduction in entropy of the posterior is computed for each test level by averaging over the two possible responses (left and right) on the next trial, weighted by the predicted probability of each response:

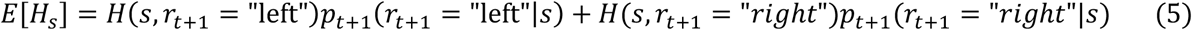

The stimulus position that minimizes the information entropy is selected for the subsequent trial 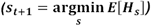 as it provides the greatest reduction in uncertainty for the α and β estimates from the current trial to the next (Shannon, 1948). That is, the stimulus position that will theoretically aid in maximally efficient parameter estimation is selected on each trial.

#### 2.5.2 Parameter estimation

Once the participant made their selection at the stimulus location, the Psi algorithm uses Bayes’ rule to compute the full posterior distribution for α and β values given the prior and the current response (r_t_):

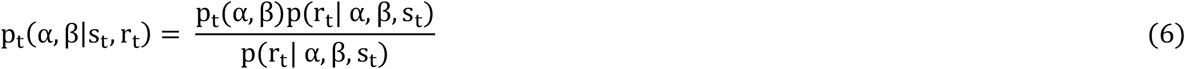

The prior, *p*_*t*_(*α, β*1, is a joint probability distribution representing the initial guess of α and β values. We selected the distributions for the priors on α and β which were the same for all participants, based on the principle of maximum entropy (Jaynes, 2003; McElreath, 2016). We assumed that the most likely α was 0 with a wide standard deviation: *α∼N*(0,201. Similarly, we assumed a reasonably wide prior distribution for β, with an expected value of 20: *β∼Exponential*(201. On subsequent trials, the posterior for trial t became the prior for trial t+1 (*p*_*t*+1_(*α, β*1 = *p*_*t*_(*α, β*|*s*_*t*_, *r*_*t*_11. The likelihood in equation 6, *p*(*r*_*t*_|*α, β, s*_*t*_1, is the probability of the participant’s response given each parameter value at the stimulus position. As suggested by Kontsevich and Tyler, we created a pair of lookup tables to improve computational efficiency, one for each response, which served as indexable likelihoods for each stimulus:

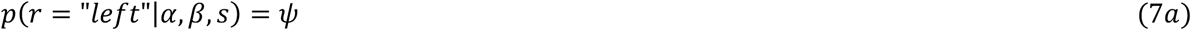

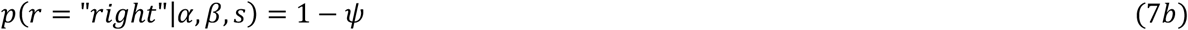

The individual α and β estimates were calculated by marginalizing over the joint posterior distribution (equation 6) and taking the mean of each marginalized distribution (Emerson, 1986). We used the final α and β estimates after the last (75^th^) trial for our agreement analyses. To determine if 75 trials was necessary, we also performed the analysis for α and β estimates after the 50th and 25th trials.

### 2.6. Statistical analysis

We used a Bayesian approach to make statistical inferences for the validity of responses and the agreement of PSE and uncertainty estimates. This approach enabled us to calculate the full posterior probability distribution of model parameters, providing a complete picture that quantifies our uncertainty. Thus, it naturally emphasizes estimation over binary decision rules in accordance with recommendations made by the American Statistical Association and others (Kruschke, 2013; Kruschke and Liddell, 2018; Wasserstein and Lazar, 2016).

For each analysis, we started by defining a statistical model that was consistent with our question and the data structure. Next, we calculated the posterior probability of all parameter values in the statistical model using Bayes’ rule, combining our prior assumptions regarding parameter values (i.e., the prior) with evidence from our data (i.e., the likelihood). We selected the distribution for each parameter’s prior as reasonably wide and uninformative, using its maximum entropy distribution (Jaynes, 2003; McElreath, 2016). We used the Pymc4 (version 4.3) library (Salvatier et al., 2016) in Python (version 3.11) to perform Markov Chain Monte Carlo (MCMC) sampling to estimate the posterior probability distributions. We drew 10,000 samples from the posterior in each of 4 chains (i.e., 40,000 total samples), using 2,000 tuning samples in each chain. We performed diagnostics for each model, ensuring parameter values were consistent across chains and checking posterior estimates for possible errors (Kruschke, 2014; McElreath, 2016). We provide the full models and code, including detailed information about the priors and diagnostics, online at https://osf.io/g8nx4/. We made inferences based on the posterior distribution of the model parameters, reporting the 95% high density interval (HDI), defined as the narrowest span of credible values that contains 95% of the posterior distribution (Kruschke, 2014). In cases in which we wanted to quantify support for the “null” hypothesis, we also calculated the percent of credible parameter values that fell within a region of practical equivalence (ROPE; Kruschke, 2014).

First, we ensured that responses were not confounded by movement direction, a concern based on a prior study (Vazquez et al., 2015). We accomplished this by determining the unique contribution of both movement directions to the probability of responding “left”, controlling for stimulus positions (which were not necessarily equal within individuals due to the adaptive nature of the Psi algorithm) and participants’ intrinsic biases towards responding “left”, using a Bayesian logistic regression model. We modeled each participant’s response data on each trial (t) as a Bernoulli distribution where the probability of responding “left” (*p*_*left*_) was impacted by the stimulus location (*X*_*stimulus*_), the movement direction (*β*_*Move direction*_), and the participant (*α*_*Participant*_):

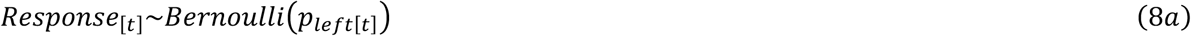

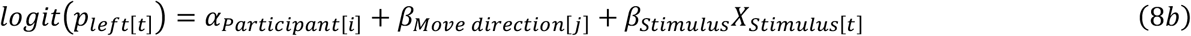

We included data from both tests for each participant in this model. Stimulus position was coded as a continuous variable. Movement direction and participant were coded as indexing variables, meaning separate *α*_*Participant*_ posteriors were computed for each participant (*i*; 13 total alphas), representing each participant’s bias to judge “left” independent of movement direction or stimulus position, and two separate group level *β*_*Move direction*_ posteriors were computed for each movement direction (*j*; forward and backward; 2 total *β*_*Move direction*_ parameters) to the stimulus position. We reasoned that if moving forward or backward had no influence on responses, the two posterior distributions for *β*_*Move direction*_ should be identical. We therefore calculated the difference in probability between the *β*_*Move direction*_ posterior distributions, setting a ROPE for this contrast between -0.5 and 0.5.

To assess test-retest agreement, we used the Bland-Altman method (Altman and Bland, 1983; Giavarina, 2015). The Bland-Altman method is recommended to assess agreement because it specifically tests for biases across the range of “true” scores of a given variable, where a proxy for the true score is the mean of an individual’s score on Test 1 and Test 2 (Bland and Altman, 1986). The method involves 3 analyses: 1) calculating the mean relative bias between the two estimates, 2) calculating the bias across “true” values, and 3) determining the limits of agreement, calculated as the mean of the difference in estimates ±1.96 times the standard deviation of the difference in estimate. To test for biases in steps 1 and 2 we applied Bayesian inference instead of the frequentist methods typically used in the Bland-Altman method. We determined if there was a mean bias by estimating the distribution of differences between Test 1 and Test 2 for the PSE and uncertainty estimates separately. We modeled these contrasts as a normal distribution, estimating the most likely μ and σ values that could have generated each individual’s (*i*) estimated difference from Test 1 to Test 2:

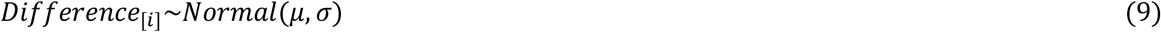

We focused our inference on the posterior distribution for μ, which, if no bias was present, should be close to zero. We therefore set a ROPE between -5 and 5 mm. We determined if there was a bias across the range of “true” values of PSE and uncertainty using a Bayesian regression analysis. The outcome variable, differences between Test 1 and Test 2 estimates for each participant (*Y*_[*i*]_), was modeled as a normal distribution with the mean (μ) being predicted by the “true” PSE/uncertainty value (*X*_*true*_):

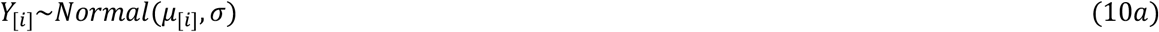

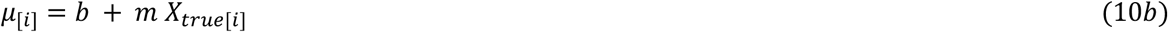

Here, a bias will manifest as an intercept (*b*) and slope (*m*) with magnitudes greater than 0. Therefore, we set a ROPE for the intercept between -5 and 5 mm and a ROPE for the slope between -0.1 and 0.1, which would translate to a 1 mm change in bias for a 10 mm change in “true” score. We defined good agreement as little to no evidence of a mean bias or a bias across true scores. Specifically, 90% of the posterior distributions calculated in the Bland-Altman analysis should not fall outside the ROPEs. We also performed a secondary analysis to determine if perfect agreement was at least plausible by characterizing the relationship between Test 1 and Test 2 for both the PSE and uncertainty estimates. Perfect agreement would result in a slope of 1 and an intercept of 0. We again used a Bayesian regression model, except the estimate on Test 2 was the outcome variable (*Y*_*TEST* 2_1 and the estimate on Test 1 was the predictor variable (*X*_*TEST* 1_):

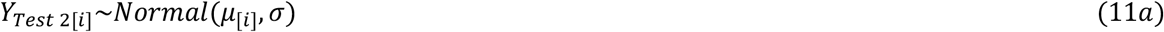

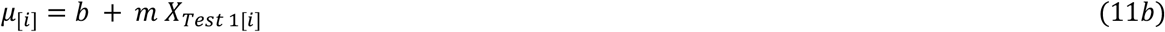

Here the HDIs for *m* characterize the relationship between Test 1 and 2.

## 3. Results and discussion

### 3.1. Movement direction did not influence participant responses

Thirteen participants (8 female, 5 male) completed the study. Data for one representative participant are displayed in Figure 1C. The psychometric functions (solid curves) were plotted from the PSE (α) and uncertainty (β) estimates after the 75^th^ trial and superimposed onto the empirical data (dots). For both tests, PSE values were positively biased such that individuals tended to feel their left and right feet were aligned when in reality their left foot was further forward of the right (Test 1 mean [95% HDI] = 12.0 mm [3.3 20.6]; Test 2 = 15.3 mm [5.3 25.2]), and uncertainty measures (i.e., SDs) were just under 20mm (Test 1 = 18.2 mm [14.4 21.8]; Test 2 = 17.5 mm [13.5 21.4]). Our first concern, based on a prior study (Vazquez et al., 2015), was to ensure that we controlled for the potential confound that movement direction may have had on responses. We determined the individual impact of movement direction on responses using a Bayesian logistic regression. If movement direction did play a major role in responses, the difference in probabilities for *β*_*Move direction* [*FORWARD*]_ vs *β*_*Move direction* [*BACKWARD*]_ would be large, making the contrast largely different from 0. However, we found that the movement direction contrast was practically equal to 0 (Figure 1D; β_Move direction [Backward]_ - β_Move direction [Forward]_ mean [95% HDI] = 0.18 [-0.07 0.43], 99.5% in ROPE). Thus, randomizing the start positions prevented biased responses.

### 3.2. Bland-Altman analysis revealed good agreement for both PSE and uncertainty estimates

We used the Bland-Altman method to assess agreement (Figure 2A). First, we calculated the limits of agreement for PSE and uncertainty estimates (−22.3 to 15.5 mm and -10.1 to 11.6 mm, respectively; dotted lines in Figure 2A). Limits of agreement for both were relatively narrow considering the scale of the measurement, and the PSE limits were consistent with the range of values observed in previous work (Vazquez et al., 2015) which used 33 more trials than the current method. Next, we characterized the mean relative bias between Test 1 and Test 2 for PSE and uncertainty estimates (histograms in Figure 2A). Both biases were very close to 0. The average bias for PSE was -3.3mm [-9.0, 2.8] (72.4% in ROPE). In other words, while the PSE was nominally lower on Test 2 than Test 1, we can be fairly confident that there is little-to-no bias for PSE measurements. With regard to uncertainty estimates, there was effectively no bias as the 95% HDI was fully within the ROPE (0.7mm [-2.8, 4.1], 98.8% in ROPE).

**Figure 2:**
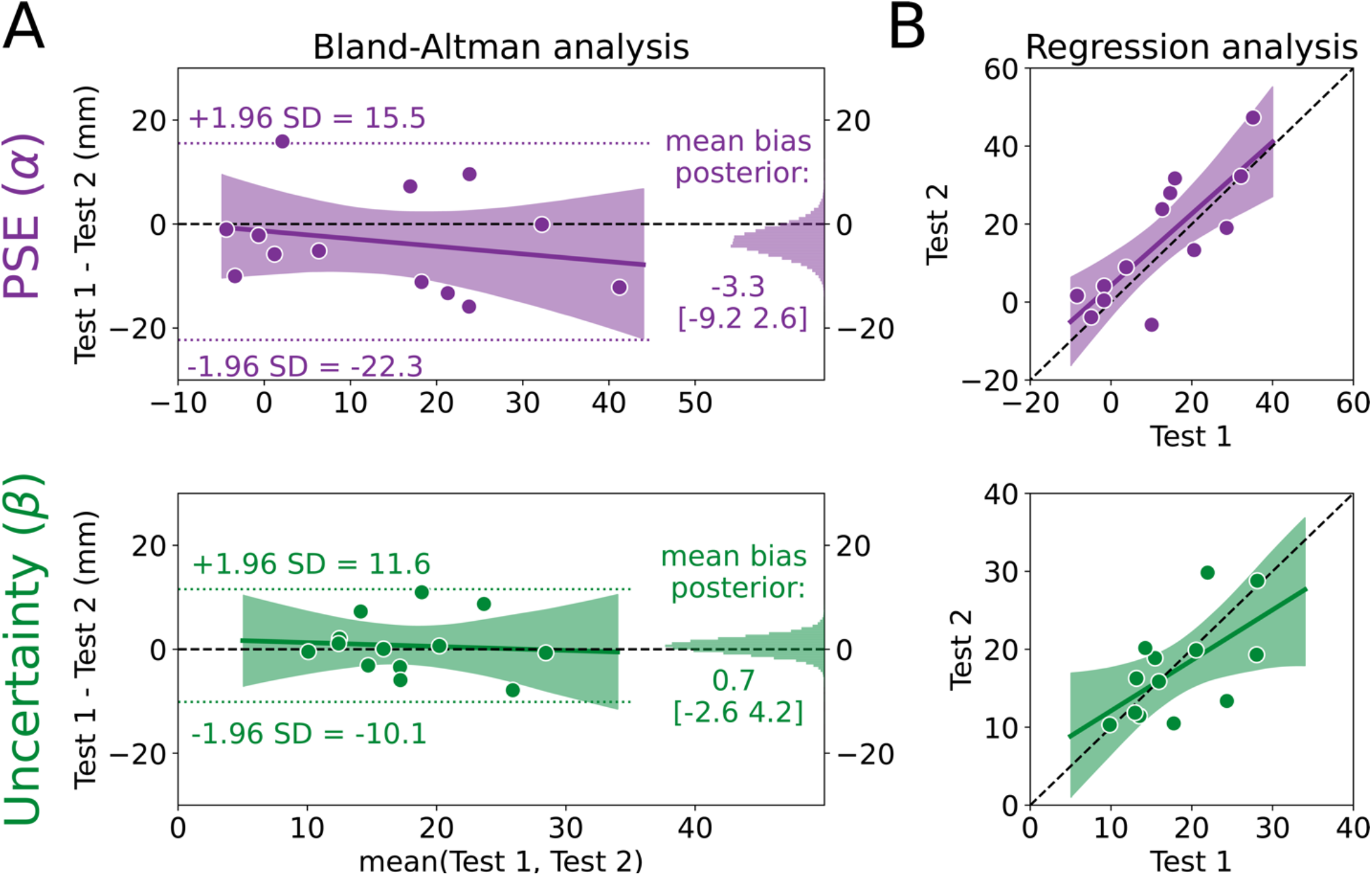
Point of subjective equality (PSE) and uncertainty estimates have good agreement. **(A)** Bland-Altman plot for the PSE (α; top) and uncertainty (β; bottom) estimates. The mean of Test 1 and Test 2 for each individual is plotted on the x-axis and the difference between the two tests is plotted on the y-axis. Thus, each dot represents one individual. The limits of agreement are plotted as the dotted lines, and the posterior distributions for the mean biases are plotted on the right sides as histograms. The most probable regression line is the solid line with the shading representing the 95% high density interval (HDI), with the black dashed line at 0 as a reference **(B)** Regression model for PSE and uncertainty. Estimates for Test 2 are plotted against estimates for Test 1 for each individual (dots) with the unity line (black dashed line) provided as a reference for perfect agreement. The most probable regression line is the solid line with the shading representing the 95% HDI.

Next, we tested for evidence of a bias across different “true” PSE and uncertainty estimates, where the true estimate is represented by the mean of an individual’s estimates on Test 1 and Test 2 (i.e., the x-axis in the Bland-Altman plot), using a Bayesian linear regression model (solid regression line and shading in Figure 2A). For PSE, there was a small negative slope, although the most credible estimates surrounded a slope estimate of zero (−0.15 [-0.56, 0.28], 29.6% in ROPE), and the intercept was unbiased (−1.3 [-9.5 6.9], 76.0% in ROPE). These results indicate that participants with larger PSE values tend have increased relative bias between test 1 and 2. However, even when considering the largest mean PSE of ∼40mm, this only produces a bias of -6mm. Therefore, we interpret this bias as negligible. For uncertainty estimates, the most probable regression was very similar to a line with 0 slope and 0 intercept (slope = -0.08 [-0.69, 0.58], 24.5% in ROPE; intercept = 2.0 [-9.8, 13.8], 58.4% in ROPE). Combined, both Bland-Altman results show that estimated PSE and uncertainty values remain largely consistent across test-retest sessions and across the range values, a sign of good agreement for this method.

As a secondary analysis, we characterized the relationship between estimates on Test 1 and Test 2 to determine if something close to perfect agreement (slope=1, intercept=0) was at least plausible, again using a Bayesian linear regression model. Whereas the Bland-Altman analysis uses information from the two tests to estimate a “true” score (the x-axes in Fig 2A), the regression analysis measures the difference between the two tests without sharing information across tests. Figure 2B shows the most credible regression lines for PSE (top) and uncertainty (bottom) estimates, with the shading representing the 95% HDI. While there were credible slope values that fell both above and below one (slope = 0.92 [0.49, 1.38], intercept = 4.2 [-3.7, 12.4]), the single best PSE estimate (maximum a posteriori) between Tests 1 and 2 was very nearly one. The uncertainty estimates indicated that this parameter is less likely to perfectly reproduce the scores on Test 1 and Test 2 (slope = 0.65 [0.09, 1.19], intercept = 5.6 [-5.1, 15.8]). Interestingly, the results of this secondary analysis seemingly conflict with the Bland-Altman analysis. However, we see them as complementary: While the average bias is low and consistent across scores, the uncertainty values are less likely to be exactly reproduced from Test 1 to Test 2 compared to the PSE values. We do not believe this is a practice effect, but rather, an instance of regression to the mean, as individuals with lower uncertainty on Test 1 demonstrated increased uncertainty on Test 2 and individuals with higher uncertainty on Test 1 demonstrated decreased uncertainty on Test 2. Consistent with reports in other psychophysical assessments, measures of uncertainty are more difficult to recover compared to PSE (King-smith and Rose, 1997; Kontsevich and Tyler, 1999; Turpin et al., 2010). Together, the results of this secondary analysis suggest that, while there is evidence of variability across test sessions, it is well within an acceptable range for psychophysical assessment. Indeed, the 95% HDI for PSE and uncertainty estimates both included what we would expect if the tests had perfect agreement.

As further evidence of good agreement in our method, and to make direct comparisons with other studies, we also calculated intraclass correlation coefficients (ICC_2,1_), another commonly used metric of agreement (Berchtold, 2016; Kottner et al., 2011; McGraw and Wong, 1996). The ICC values for PSE (ICC_2,1_ [95% CI] = 0.80 [0.48, 0.93]) and uncertainty (0.61 [0.05, 0.85]) indicate good agreement (Portney and Watkins, 2009). These values fall within ranges of other psychophysics studies using the Psi algorithm and the method of constants, as those studies report ICCs for PSE estimates that range between 0.48 to 0.96 for the Psi algorithm (Schilling et al., 2017; Silva et al., 2020), and between 0.77 to 0.88 for the method of constants (Nicholson et al., 1997). Our method also compares favorably to other proprioception-specific assessments with ICCs ranging from 0.11 to 0.95 for measures of proprioceptive accuracy (Antcliff et al., 2021; Arvin et al., 2015; Deshpande et al., 2003; Gorst et al., 2020; Hillier et al., 2015; Rinderknecht et al., 2018), and from 0.0 to 0.64 for measures of proprioceptive variability (Juul-Kristensen et al., 2008; Rahlf et al., 2019; Strong et al., 2021). The fact that the current method has good agreement for both proprioceptive PSE and uncertainty opens the door for future studies to assess the importance of uncertainty to lower limb function. We provide a more detailed account of these studies in a table available online at our osf page.

### 3.3. Agreement was similar after 50 trials

One of our primary goals for this method was to maximize efficiency. Participants took an average of 20±1 minutes to complete testing (including the two short walking breaks). The simplest way to reduce this time would be to reduce the number of trials, however, there is a risk that reducing the number of trials would also reduce the agreement of the PSE and uncertainty estimates. Since the Psi algorithm estimates PSE and uncertainty after every trial, we can assess agreement after any trial during the test. As a post-hoc analysis, we chose to assess agreement after 50 trials as the test time at this point would have been between 10-15 minutes. We found that all our measures of agreement were similar after 50 and 75 trials (Table 1). In contrast, examining agreement after 25 trials revealed a substantial reduction in agreement. Therefore, decreasing the number of trials to 50 would significantly reduce the time of the test without sacrificing agreement, providing a reasonable timeframe for a test to be placed in the middle of a motor learning paradigm or to be used for a balance and falls risk assessment.

**Table 1:** Agreement comparison after 50 and 75 trials.

| | Limits of agreement (mm) | Bland Altman Mean Bias (mm; % in ROPE) | Bland Altman Bias Regression (mm; % in ROPE) | Test 1 vs Test 2 Regression | $ICC_{2,1}$ [95% CI] |
| --- | --- | --- | --- | --- | --- |
| **PSE ($\alpha$)** |  |  |  |  |  |
| 50 trials | -22.3 to 15.7 | -3.2 [-9.2 2.6] (73.5%) | Slope = -0.17 [-0.61 0.29] (26.2%)<br>Intercept = -1.1 [-9.3 7.1] (77.0%) | Slope= 0.91 [0.45 1.40]<br>Intercept = 4.1 [-3.9 12.0] | 0.78 [0.43, 0.92] |
| 75 trials | -22.3 to 15.5 | -3.3 [-9.0 2.8] (72.4%) | Slope = -0.15 [-0.56 0.28] (29.6%)<br>Intercept = -1.3 [-9.5 6.9] (76.0%) | Slope= 0.92 [0.49 1.38]<br>Intercept = 4.2 [-3.7 12.4] | 0.80 [0.48, 0.93] |
| **Uncertainty ($\beta$)** |  |  |  |  |  |
| 50 trials | -9.3 to 10.2 | 0.5 [-2.7 3.5] (99.5%) | Slope = 0.03 [-0.65 0.69] (24.3%)<br>Intercept = 0.0 [-12.5 12.2] (60.0%) | Slope= 0.57 [0.03 1.09]<br>Intercept = 7.4 [-2.8 17.4] | 0.58 [0.05, 0.85] |
| 75 trials | -10.1 to 11.6 | 0.7 [-2.8 4.1] (98.8%) | Slope = -0.08 [-0.69 0.58] (24.5%)<br>Intercept = 2.0 [-9.8 13.8] (58.4%) | Slope= 0.65 [0.09 1.19]<br>Intercept = 5.6 [-5.6 15.8] | 0.61 [0.05, 0.86] |

**Table 1: Agreement comparison after 50 and 75 trials**
|  | Limits of agreement (mm) | Bland Altman Mean Bias (mm; % in ROPE) | Bland Altman Bias Regression (mm; % in ROPE) | Test 1 vs Test 2 Regression | ICC<sub>2,1</sub> [95% CI] |
| --- | --- | --- | --- | --- | --- |
| **PSE (α)** |  |  |  |  |  |
| 50 trials | -22.3 to 15.7 | -3.2 [-9.2 2.6] (73.5%) | Slope = -0.17 [-0.61 0.29] (26.2%)<br>Intercept = -1.1 [-9.3 7.1] (77.0%) | Slope= 0.91 [0.45 1.40]<br>Intercept = 4.1 [-3.9 12.0] | 0.78<br>[0.43, 0.92] |
| 75 trials | -22.3 to 15.5 | -3.3 [-9.0 2.8] (72.4%) | Slope = -0.15 [-0.56 0.28] (29.6%)<br>Intercept = -1.3 [-9.5 6.9] (76.0%) | Slope= 0.92 [0.49 1.38]<br>Intercept = 4.2 [-3.7 12.4] | 0.80<br>[0.48, 0.93] |
| **Uncertainty (β)** |  |  |  |  |  |
| 50 trials | -9.3 to 10.2 | 0.5 [-2.7 3.5] (99.5%) | Slope = 0.03 [-0.65 0.69] (24.3%)<br>Intercept = 0.0 [-12.5 12.2] (60.0%) | Slope= 0.57 [0.03 1.09]<br>Intercept = 7.4 [-2.8 17.4] | 0.58<br>[0.05, 0.85] |
| 75 trials | -10.1 to 11.6 | 0.7 [-2.8 4.1] (98.8%) | Slope = -0.08 [-0.69 0.58] (24.5%)<br>Intercept = 2.0 [-9.8 13.8] (58.4%) | Slope= 0.65 [0.09 1.19]<br>Intercept = 5.6 [-5.6 15.8] | 0.61<br>[0.05, 0.86] |

### 3.4. Limitations and future directions

Despite the efficiency and good agreement of this method, we also recognize some of its limitations. For instance, one way to increase ecological validity when measuring proprioception would be to assess three-dimensional position sense as opposed to measuring proprioception in only the sagittal plane as we have done here. However, we believe our results should at least generalize to locomotor adaptation on the split-belt, given that, in this context, adaptation is often measured in sagittal plane kinematics like step length (Reisman et al., 2005). Another potential limitation is that during standing, although participants’ limbs were passively moved by the motorized treadmill, it is impossible to avoid some subtle movements and associated muscle activity. While no overt movements were observed during testing, in theory, any voluntary muscle activity might provide additional information about where the limb is located due to an efference copy of the motor command (Wolpert et al., 1995). However, in our case, such a compromise was unavoidable, as the primary goal of this study was to develop an assessment of lower limb position sense in a context that most closely resembles gait.

In addition to measuring proprioception in standing, we assessed position sense of the whole lower limb by asking individuals to focus on the end effector (foot position) when making their judgements. Interestingly, there are few static lower extremity proprioception studies that ask individuals to focus on end effector position (but see Sigmundsson et al., 2000; Vazquez et al., 2015), despite the functional significance of foot position to successful gait (e.g., stepping over curbs, trail running, etc.). Conversely, upper extremity studies frequently assess whole limb proprioception by having participants make judgments regarding end effector position (hand or finger (Jones et al., 2010; Vindras et al., 1998). These methods have characterized the precision of limb position sense (van Beers et al., 1998), the relationship between static proprioception and voluntary reaching movements (Jones et al., 2010), and the importance of static proprioception to sensorimotor adaptation (Clayton et al., 2014; Cressman and Henriques, 2010; Simani et al., 2007; Tsay et al., 2021; van Beers et al., 2002). Unfortunately, similar lines of research are absent in locomotion, despite the critical role of proprioception in walking (Hiebert et al., 1996; Kriellaars et al., 1994; Pearson, 2004; Roden-Reynolds et al., 2015; Whelan et al., 1995). We speculate this is due at least in part to the previous absence of reliable and efficient methods of assessing lower limb proprioception in an upright, functional, and multi-joint context. We hope the current method opens the door to increasing understanding of the relationship between lower limb proprioception and locomotor adaptation in young, neurotypical adults as well as in older and neurologic populations, for whom changes in proprioception may have a significant impact on their ability to adapt gait patterns (Bruijn et al., 2012; Bunday and Bronstein, 2009; Lam and Pearson, 2002; Pearson, 2000; Santuz et al., 2022).

Efficient and reliable, whole lower limb proprioceptive measurements during standing should be useful for non-adaptation studies as well. For example, proprioception is an important part of multifactorial falls assessments (Lord et al., 1991), but the proprioception test previously recommended involves active toe position matching in a non-weight bearing position. The current method offers a more ecologically valid lower limb proprioception test since it is much closer in context to when falls most often occur: during weight bearing activities like walking and transfers (Talbot et al., 2005). Furthermore, since most lower limb proprioception tests only measure proprioceptive accuracy or bias (Han et al., 2016; Hillier et al., 2015; Horváth et al., 2022), it is unknown how proprioceptive uncertainty may relate to falls, something that can now be empirically assessed using the current method.

As there are very few studies reporting PSE and uncertainty values for the lower limbs, we were interested in comparing our values to those reported in upper extremity studies. We found that the average uncertainty estimates in the present study were quite similar to values found in upper extremity proprioception studies (Jones et al., 2010), with both falling just under 2 cm. Additionally, we found that PSE estimates here were biased, a finding consistent with intrinsic biases in perceived hand/arm position (Fuentes and Bastian, 2010; Ingram et al., 2019; Jones et al., 2010; van Beers et al., 1996). Interestingly, based on a small pilot study we conducted, this bias may have been related to which limb was moved during testing, as we observed a negative PSE in participants whose right leg was moved (n=4). However, given the small sample size and the high uncertainty in the estimates, further testing is required to determine the extent to which movement of the foot contributes to estimated biases. Importantly, biased estimates in upper extremity studies have been linked to biases in reaching direction (Jones et al., 2010; Vindras et al., 1998), suggesting that future work in the lower extremities may benefit from examining whether PSE biases are related to step length or other gait-related movements.

## 4. Conclusion

We developed and tested the reliability of a lower limb proprioception assessment in a gait-specific context. This method is efficient, requiring only 50 trials to reliably estimate both the PSE and uncertainty of lower limb proprioception. We believe this method will aid future mechanistic understanding of locomotor adaptation and serve as a useful tool for basic and clinical researchers studying balance and falls.

## Acknowledgements

This work was supported by the National Institutes of Health [K12-HD055931 (HEK), R01-AG071585 (SMM), S10-RR028114 (SMM)], the National Science Foundation [M3X 1934650 (HEK)], and partial funding from the University of Delaware Graduate College (JMW). We want to extend a special thank you to Saunders Penn for his help creating Figure 1A and B and for his help performing data collections.

